# Engineered hydrogel biomaterials facilitate lung progenitor cell differentiation from induced pluripotent stem cells

**DOI:** 10.1101/2024.12.16.628773

**Authors:** Alicia E. Tanneberger, Rachel Blomberg, Ganna Bilousova, Amy L. Ryan, Chelsea M. Magin

**Affiliations:** Department of Bioengineering, University of Colorado, Denver | Anschutz Medical Campus, Aurora, CO, USA; Department of Dermatology, Gates Institute, University of Colorado, Anschutz Medical Campus, Aurora, CO, USA; Department of Anatomy and Cell Biology, Carver College of Medicine, University of Iowa, Iowa City, IA, USA; Department of Pediatrics, University of Colorado, Denver | Anschutz Medical Campus, Aurora, CO, USA; Division of Pulmonary Sciences and Critical Care Medicine, Department of Medicine, University of Colorado, Anschutz Medical Campus, Aurora, CO, USA

**Keywords:** Lung progenitor cells, induced pluripotent stem cells, hydrogels, differentiation, biomaterials

## Abstract

Lung progenitor (LP) cells identified by the expression of transcription factor NK2 homeobox 1 (NKX2.1) are essential for development of all lung epithelial cell types and hold tremendous potential for pulmonary research and translational regenerative medicine applications. Here we present engineered hydrogels as a promising alternative to the naturally derived materials that are often used to differentiate human induced pluripotent stem cells (iPSCs) into LP cells. Poly(ethylene glycol) norbornene (PEGNB) hydrogels with defined composition were used to systematically investigate the role of microenvironmental stiffness, cell origin, and splitting during the differentiation process. Results demonstrated each factor impacted LP differentiation efficiency. Soft hydrogels replicating healthy lung stiffness (Elastic modulus (E), E = 4.00 ± 0.25 kPa) produced the highest proportion of LP cells (54% by flow cytometry), stiff hydrogels (E = 18.83 ± 2.24 kPa) resulted in 48% differentiation efficiency, and a thin coating of Matrigel on tissue culture (TC) plastic (E∼3 GPa) resulted in the lowest proportion of LP cells (32%) at the end of the non-split differentiation protocol. Collectively these results showed that engineered hydrogels enabled control over parameters that impacted differentiation and produced LP cells using well-defined microenvironments that may improve our ability to translate iPSC-derived LP cells into clinical applications.

**NEW & NOTEWORTHY:** Standard iPSC differentiation protocols rely on Matrigel, a basement membrane extract from mouse sarcoma cells that is poorly defined and exhibits significant batch-to-batch variation. Due to these limitations Matrigel-derived products have never been approved by the Food and Drug Administration. This study introduces a novel method for differentiating iPSCs into lung progenitor cells using well-defined hydrogel substrates. These biomaterials not only enhance differentiation efficiency, but also streamline the regulatory pathway, facilitating their potential therapeutic application.

## INTRODUCTION

During development, lung progenitor (LP) cells are the precursor to all airway and alveolar epithelial cells (1, 2). In the mature lung LP cells have differentiated into basal stem cells and alveolar type 2 cells, the resident stem cells responsible for regeneration and injury repair (3). Although there are broadly acknowledged advantages to culturing epithelial stem cells *ex vivo*, it has been difficult to produce a pure mature cell population that retains the desired phenotype after isolation from human lungs (2, 4–6). For this reason, the generation of LP cells from human pluripotent stem cells (PSCs) including both embryonic stem cells (ECSs) (7–9) and induced pluripotent stem cells (iPSCs) (10–14) is an attractive alternative with the potential to generate sufficient cells for applications such as autologous cell therapy. Despite subtle variations in differentiation protocols, all approaches require cells to progress through the definitive endoderm (DE) and anterior foregut endoderm (AFE) phases before LP induction, mimicking the core stages of embryonic lung development (9, 13–15). The transcription factor NK2 homeobox 1 (NKX2.1) is one of the most widely used LP cell markers. Extended differentiation protocols have shown that NKX2.1+ cells can give rise to both proximal airway (9, 10) and distal lung epithelial cells (9, 13, 14). High expression of carboxypeptidase M (CPM^hi^), a cell surface marker that correlates well with NKX2.1 levels, has enabled cell sorting to isolate LP cells (9, 13, 14, 16).

Differentiation efficiencies between 5-90% are considered successful in established protocols (9, 13, 14, 17). There is clearly an unmet need to reduce this variability and enhance average differentiation efficacies. A common feature of PSC differentiation protocols that may contribute to this high level of variation is the almost exclusive reliance on animal-derived materials, such as Matrigel. Matrigel is a basement membrane extract from mouse sarcoma, a tumor rich in many extracellular matrix (ECM) proteins including collagen IV, laminin, proteoglycans, entactin, and numerous growth factors (18). Mechanically, the elastic modulus (E) of 3D Matrigel hydrogels measures approximately 0.05 kPa (19), which falls well below the range of healthy lung tissue (E=1-5 kPa) (20–22). Furthermore, when low concentrations of Matrigel are coated on tissue culture (TC) plastic surfaces, cells are exposed to the supraphysiological mechanical stiffness of the TC plastic (E∼3 GPa) (23–25). For these reasons, this naturally derived material is not well-suited to the specific changes, testing, and iterative improvements needed to refine it for specific cell culture applications. Likewise, due to the poorly defined composition of Matrigel and significant batch-to-batch variations, it is difficult to obtain approval for using cells cultured with Matrigel in clinical trials. In fact no Matrigel-derived products have been approved by the Food and Drug Administration (26).

Researchers have begun exploring engineered biomaterials as alternative cell culture microenvironments to address these limitations (27–30). Loebel et al. differentiated iPSC-derived LP cells into induced alveolar epithelial type 2 (iAT2) cells using Matrigel, then eliminated Matrigel while propagating iAT2 cells within hyaluronic acid-based hydrogels (31). Engineered hydrogel biomaterials such as those containing poly(ethylene glycol) norbornene (PEGNB) can be reliably reproduced and uniquely tailored to control the composition and mechanical properties (20). Adjusting the weight percent (wt%) of PEGNB in a hydrogel formulation can produce biomaterials that mimic a wide range of physiologically relevant lung stiffnesses. This versatile backbone can also be combined with a wide range of crosslinkers to create cell-instructive microenvironments. For example, peptide sequences degradable by cell secreted matrix metalloproteinase (MMP) enzymes can be incorporated to enable hydrogel remodeling (32–34). Cell adhesive peptides that mimic binding sites on proteins like fibronectin (CGRGDS) or laminin (CGYIGSR) can also be added into the hydrogel formulation to enhance cell binding (35–37). These modifications are well-defined and highly controlled.

The goal of the present study was to eliminate the use of naturally derived materials during iPSC differentiation. Here, we present a systematic study that investigates the impact of substrate elastic modulus, cell line, and splitting as iPSCs are differentiated into LP cells on 2D engineered hydrogels. Quantification of NKX2.1 protein and gene expression showed that engineered hydrogels promoted similar and/or improved differentiation efficiencies compared to traditional Matrigel-based differentiation protocols (13, 14). Flow cytometric analysis demonstrated an increase in the number of CPM^hi^ expressing cells when non-split iPSCs were differentiated on soft PEGNB hydrogels. This technological advancement demonstrates the feasibility of differentiating iPSCs into LP cells under Matrigel-free conditions and has the potential to support the development of protocols that enhance the acceptance of iPSC-derived cells in clinical trials.

## MATERIALS & METHODS

Methods are described in detail in the supplemental documents.

### Maintenance of human induced pluripotent stem cell lines

Three iPSC (Supplemental Table S1) lines were expanded and maintained for at least two passages before differentiation.

### 2D hydrogel synthesis and characterization

A poly(ethylene glycol)-norbornene (PEGNB; 8-arm, 10 kg/mol) macromer was synthesized and characterized as previously described (38–40). PEGNB hydrogel pre-cursor solutions were tailored to produce either soft (5.3 wt% PEGNB) or stiff (7.75 wt% PEGNB) that were crosslinked with an MMP9-degradable peptide sequence, decorated with 2 mM cell-adhesive peptides (CGRGDS and CGYIGSR), and contained 2 mg/mL entrapped laminin/entactin protein complex. Hydrogel elastic modulus (E) was measured using parallel-plate rheology, as previously described (20, 40). Hydrogels for differentiation experiments were covalently linked to glass coverslips during polymerization, swollen in mTeSR Plus medium (STEMCELL Technologies, cat. # 100-0276) and maintained at 37℃ and 5% CO2 prior to cell seeding.

### iPSC-to-LP cell differentiation

When undifferentiated iPSCs reached approximately 80% confluency (Day 0), the cells were dissociated from TC plastic flasks and seeded at 200,000 cells/cm^2^ onto either Matrigel-coated TC plastic, soft hydrogels, or stiff hydrogels. On day 0, iPSCs were maintained in mTeSR Plus medium supplemented with 10 µM Y-27632 dihydrochloride. On day 1, differentiation was initiated using a DE kit (StemDiff) following the manufacturer’s instructions. On day 5, a subset of samples was split at a 1:3 ratio (DE-split) and cultured overnight in DS/SB medium (13, 14) supplemented with 10 µM Y-27632 dihydrochloride, a Rho kinase (ROCK) inhibitor. Non-split samples were maintained in the same medium. On day 6, the medium for all samples were replaced with DS/SB medium (13, 14). On day 8, all samples were switched to CBRa medium (13, 14), which was replenished every 48 h until day 18.

### Immunostaining and image analysis

On day 18, cells were removed from culture substrates and deposited onto glass slides. Samples were stained for NKX2.1 and counterstained with DAPI. Negative controls were prepared identically but omitting the primary antibody. Imaging was performed using an Olympus BX63 upright microscope. Six random areas per slide were imaged under a 4x objective. Images were post-processed to remove background fluorescence in ImageJ Software (NIH) and quantified using the StarDist plugin to measure the percentage of NKX2.1-positive cells.

### RNA isolation, cDNA synthesis, and gene expression analysis

On day 18, RNA was isolated from samples following a TRIzol (Fisher Scientific, cat. #15-596-026) isolation protocol. The isolated RNA was converted into cDNA using a high-capacity cDNA Reverse Transcription Kit (Applied Biosystems, cat. #4368814). Reverse transcription-quantitative polymerase chain reaction (RT-qPCR) assessed gene expression of NKX2.1. All gene expression data were normalized using a natural log transformation before relative gene expression was calculated using a 2^!Δ#$^ approach. Following statistical analyses values were untransformed by exponentiating the natural log-transformed data and subsequently presented in the figures.

### Flow cytometry

Cells collected at the end of LP induction (day 18) were collected, passed through a 70 µM strainer, and resuspended in flow cytometry (FC) buffer containing 2% fetal bovine serum (FBS) in phosphate buffered saline (PBS). After counting, cells were stained with a mouse anti-human carboxypeptidase M (CPM, Fujifilm Wako Chemicals, cat. #014-27501) and a rat anti-mouse PE-Cyanine7 secondary antibody (PE-Cy7, Thermo Scientific, cat. # 25-4015-82) to assess CPM^hi^ expression. Calcein Violet-AM (Fisher Scientific, cat. #501145046) assessed viability. Flow cytometric analysis was performed using a CYTEK Aurora 5 laser spectral cytometer.

### Statistical Analysis

For all immunofluorescence (IF) staining and RT-qPCR quantification, results included three biological replicates with six to eight technical replicates. For flow cytometry quantification, results included three biological replicates, each with three to four technical replicates. The IF and flow datasets were assessed for normality using a Shapiro-Wilk test and passed. RT-qPCR results were normalized using natural log transformation. Two-way ANOVA and a Tukey test for multiple comparisons assessed statistical differences within the same type of splitting condition (non-split or DE-split) for IF, gene expression, and flow datasets (GraphPad Prism). To identify the best differentiation conditions, a custom design of experiments (DOE) was generated with JMP software (SAS). A standard least squares analysis and a three-way ANOVA with all interactions determined the main and interaction effects driving iPSC-to-LP differentiation.

## RESULTS

### PEGNB Hydrogels Supported iPSC Differentiation

Hydrogels comprised of a norbornene-conjugated, eight-arm 10 kg/mol PEG macromer backbone, an MMP9-degradable crosslinker, two integrin-binding peptide mimics, and entrapped laminin/entactin (Fig. 1A) facilitated iPSC adhesion. To assess the efficacy of iPSC differentiation into LP cells on soft and stiff hydrogels, results were compared to a standard differentiation protocol that involves culturing iPSCs on Matrigel-coated TC plastic (Fig. 1A) (8–10, 13, 14). The two discrete stiffness hydrogels, soft and stiff, were designed to mimic the stiffness of human lung tissue (20, 21, 41), while the Matrigel coated TC-plastic plates served as a third substrate with supraphysiological stiffness (23, 24). Soft hydrogels achieved an average elastic modulus value of 4.00 ± 0.25 kPa, which falls within the range of healthy lung tissue (1-5 kPa) (20–22, 41) (Fig. 1B). Stiff hydrogels exhibited an average elastic modulus value of 18.83 ± 2.24 kPa, which is more representative of injured or diseased lung tissue (20–22, 41) (Fig. 1B).

**Figure 1.**
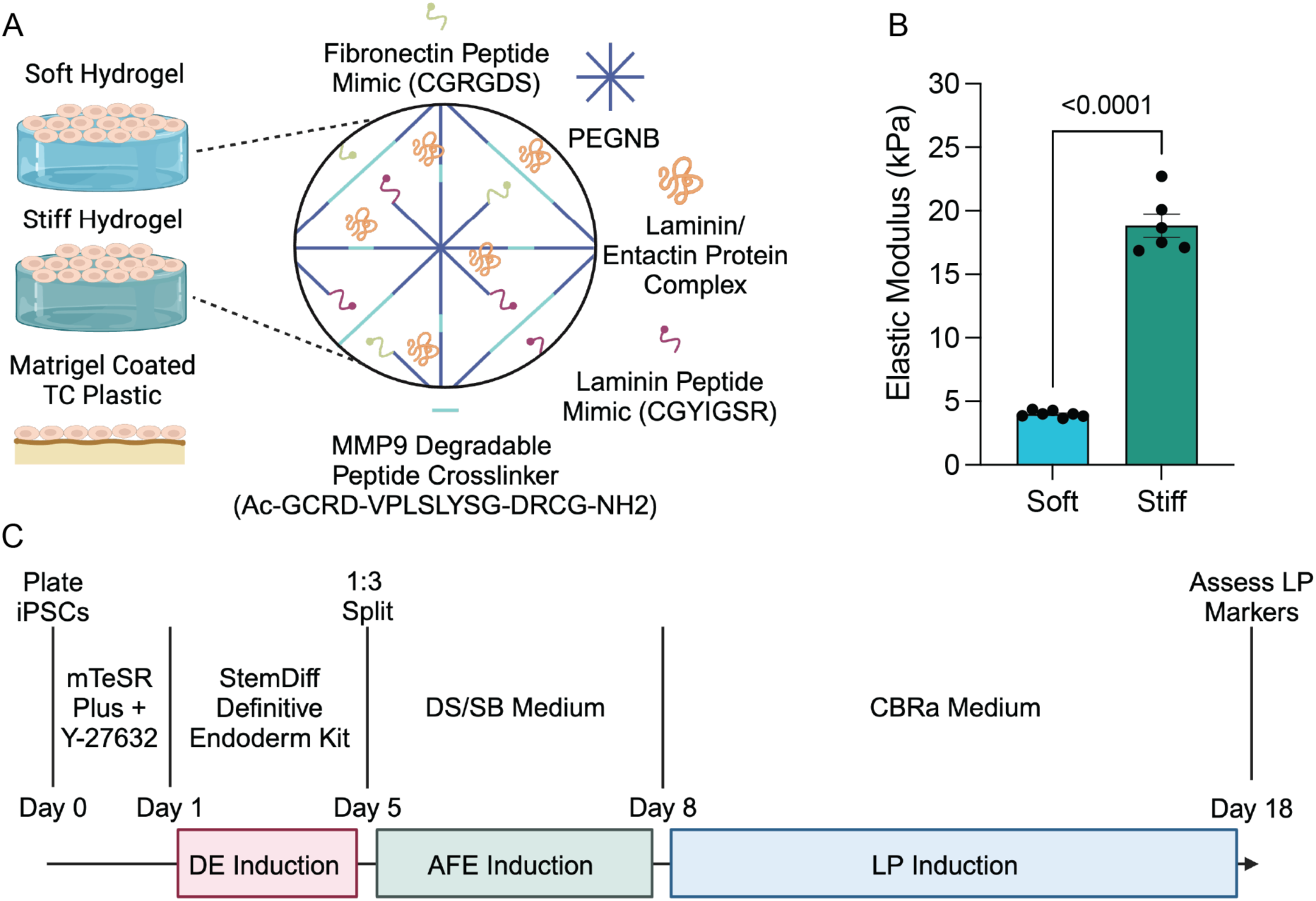
Engineered PEGNB hydrogels supported iPSC differentiation into LP cells. A) Schematic of hydrogel formulations that contained PEGNB, MMP9-degradable crosslinker, CGYIGSR, CGRGDS, and entrapped laminin/entactin. B) Rheological measurements for the average elastic modulus (E) of soft hydrogels (E = 4.00 ± 0.25 kPa, n=7) and stiff hydrogels (E = 18.83 ± 2.24 kPa, n=6). Columns represent mean ± SEM. Each dot represents an individual hydrogel sample. Statistical significance was determined by an unpaired t-test. C) Schematic representation of the differentiation protocol timeline.

Engineered PEGNB hydrogels provided iPSCs with controlled and physiologically relevant biochemical and mechanical cues throughout the 18-day differentiation protocol (13, 14) (Fig. 1C). First, undifferentiated iPSCs were seeded onto either soft hydrogels, stiff hydrogels, or Matrigel-coated TC plastic and maintained in mTeSR Plus medium supplemented with a ROCK inhibitor for 24 h. Next, DE differentiation was initiated on days 1-4 using a commercially available kit (9, 13, 14, 42). On day 5, half of the samples were split at a 1:3 ratio, and all samples were switched to DS/SB medium (13, 14, 17) promoted differentiation into AFE cells over the next 72 h. Starting on day 8, samples were transitioned into CBRa medium (13, 14, 17) and maintained for 10 days to induce LP differentiation. On day 18, cells were assessed for LP markers through IF staining, gene expression, and flow cytometry. This protocol was repeated with three individual iPSC lines (Supplemental Table S1).

### NKX2.1 Protein and Gene Expression Confirmed LP Cell Differentiation

At the end of the differentiation protocol, cells were assessed for NKX2.1 protein and mRNA expression. Immunostaining for nuclear NKX2.1 protein indicated specific expression of this well-established LP marker across all sample conditions (Fig. 2A) with no non-specific secondary antibody staining on negative controls (Supplemental Fig. S1). The average percentage of NKX2.1+ expressing cells was higher in all the non-split conditions, compared to DE-split samples. For instance, within the Matrigel on TC plastic samples, 46.39% ± 3.75% of cells expressed NKX2.1 protein in the non-split condition, versus 32.77% ± 4.47% in the DE- split condition (Fig. 2B). Similarly, within the stiff hydrogel samples, 48.19% ± 4.36% of cells expressed NKX2.1 protein in the non-split condition, compared to 26.45% ± 3.49% in the DE- split condition (Fig. 2B). This trend was also observed within the soft hydrogel condition where 42.27% ± 5.10% of cells expressed NKX2.1 protein in the non-split condition, versus 31.29% ± 3.92% in the DE-split condition (Fig. 2B). Overall, the lack of statistically significant changes in the number of cells expressing NKX2.1 protein by IF between the engineered hydrogel conditions and the standard Matrigel on TC plastic standard indicates that engineered hydrogels produced similar results while providing a well-defined and tunable cell culture substrate.

**Figure 2.**
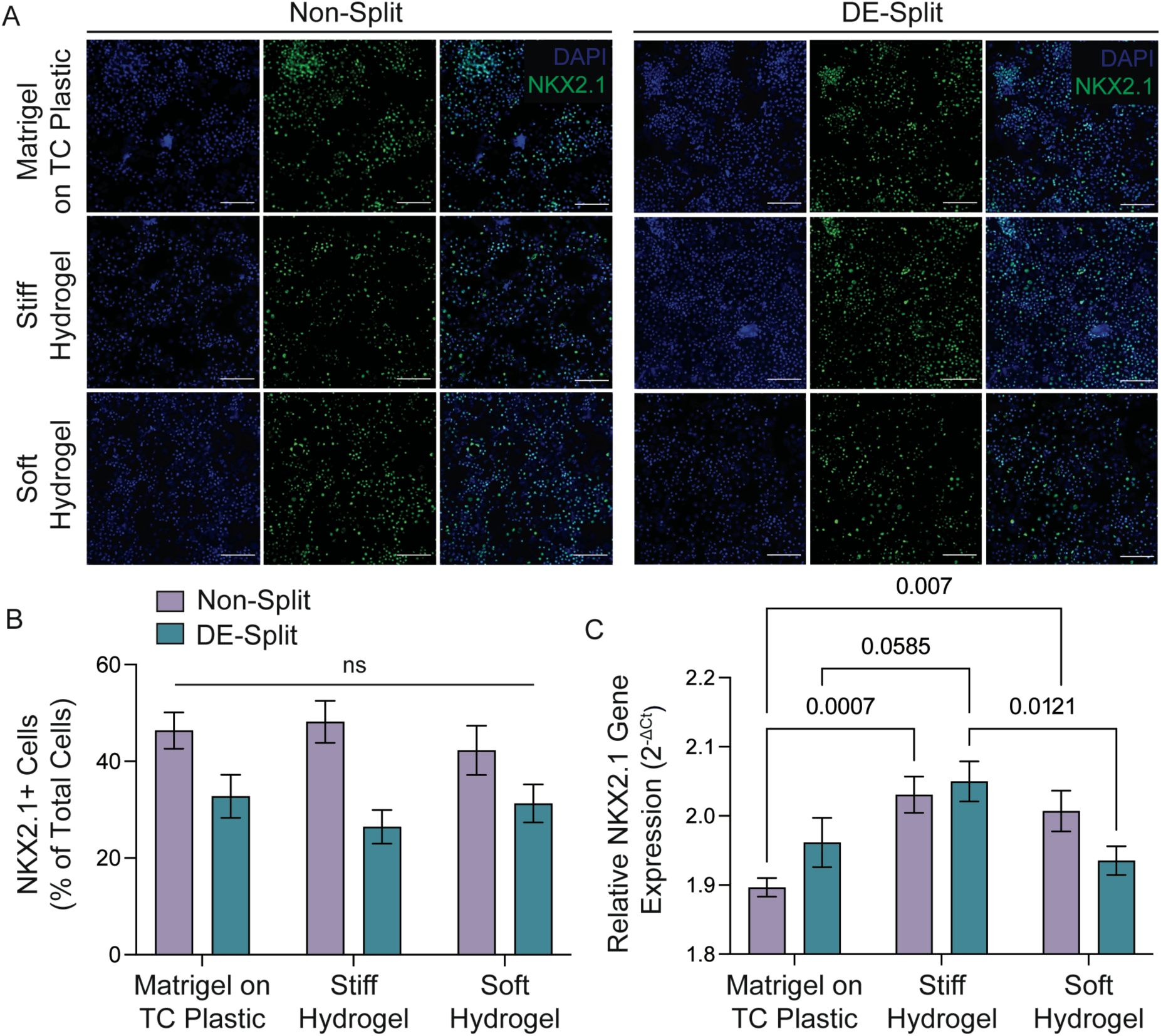
Differentiating iPSCs into LP cells on engineered PEGNB hydrogels yielded IF and gene expression results with differentiation efficiencies comparable to or higher than traditional Matrigel-based protocols. A) Representative day 18 IF images showed robust NKX2.1 expression across all non-split and DE-split substrate conditions (NKX2.1: green, DAPI: blue, scale bars = 100 μm). B) Quantification of NKX2.1-positive cells showed no statistically significant (ns) differences across substrates and splitting groups (Two-way ANOVA, Tukey Test). Columns represent mean ± SEM, n=18. C) Quantification of relative NKX2.1 gene expression showed statistically significant increases in LP differentiation on the soft and stiff hydrogel conditions compared to Matrigel on TC plastic within each splitting group (Two-way ANOVA, Tukey Test). Columns represent mean ± SEM, n=18-22.

Relative NKX2.1 gene expression was also measured, and different trends emerged. Stiff hydrogels increased LP cell differentiation efficiency by 1.07-fold compared to Matrigel on TC plastic in the non-split condition (p=0.0007) and by 1.05-fold in the DE-split condition (p=0.0585) (Fig. 2C). Similarly soft hydrogels increased differentiation efficiency by 1.05-fold compared to Matrigel on TC plastic in the non-split condition (p=0.007) (Fig. 2C). There was a decrease in NKX2.1 relative gene expression in the soft DE-split samples when compared to the stiff DE-split samples (p=0.0121) (Fig. 2C). These results suggest the impact of substrate stiffness was more apparent at the gene expression level than through immunostaining.

### Flow Cytometry Results Showed Differentiating iPSCs on Tunable Stiffness Hydrogels Increased LP Cell Differentiation Efficiency

While both IF staining and gene expression allowed for NKX2.1 end-point assessment, these techniques do not enable sorting and purification of the LP cell subpopulation. Hence, we sought to profile the iPSC-derived cells for a known LP cell surface marker, CPM^hi^, to demonstrate how this protocol could be continued to facilitate purification and further differentiation of iPSCs at the end of the LP induction. After preparing the necessary controls and staining the samples for CPM, flow cytometric analysis was performed to provide a third modality of LP verification and validate the use of engineered hydrogels in this differentiation protocol. A flow gating approach was designed and applied across all sample conditions and iPSC lines.

Debris was first removed using forward scatter area (FSC-A) versus side scatter area (SSC-A) (Fig. 3A). Then, an FSC-A versus FSC height (FSC-H) gate (Fig. 3A) was applied to remove doublets. Calcein Violet-AM dye was used to stain live cells through intracellular esterase activity and PE-Cy7 was used to measure CPM expression. Therefore, gates for FSC-A versus live cells and FSC-A versus CPM^hi^ expression were applied (Fig. 3A) and enabled quantification (Fig. 3B). While this standardized gating approach was used across all biological replicates, gates were infrequently and slightly adjusted as needed between technical replicates, taking into consideration the compensation, unstained, and secondary only controls for that run.

**Figure 3.**
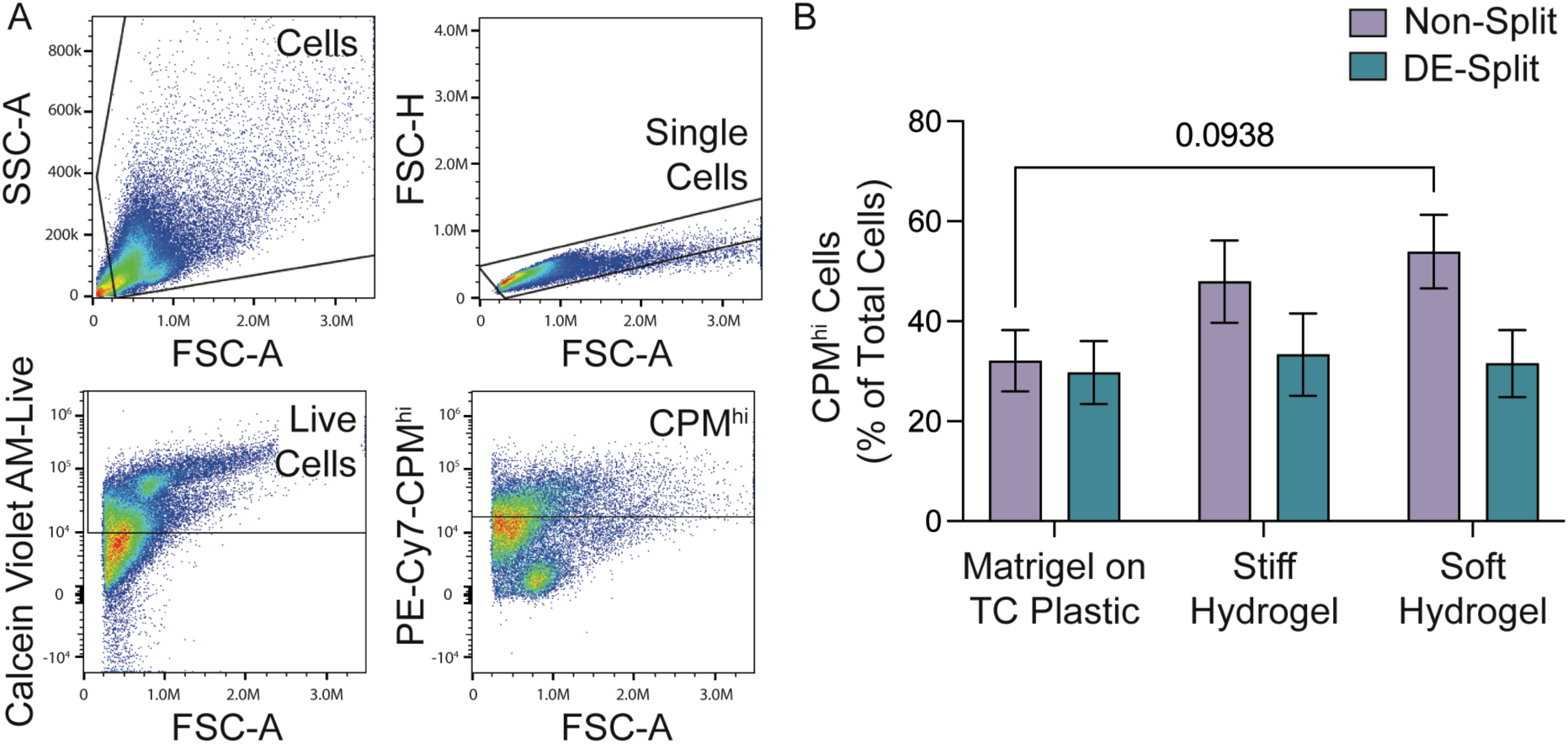
Soft hydrogels in the non-split condition showed the highest percentage of CPM^hi^ expression cells by flow cytometry. A) Representative images of the flow cytometry gating strategy that was applied to exclude debris, doublets, and dead cells to measure the percentage of CPM^hi^ expressing cells at the end of the differentiation protocol (FSC-A: forward scatter area, SSC-A: side scatter area, FSC-H: forward scatter height). B) Quantification of the percentage of CPM^hi^ expressing cells showed higher expression in the non-split soft hydrogel condition compared to Matrigel on TC plastic. Within the soft hydrogel conditions, there was also a significant decrease in CPM^hi^ expression when the cells were split at the DE stage. Columns represent mean ± SEM, n=9-10. Statistical significance was determined by a two-way ANOVA and a Tukey test within each type of splitting dataset. Direct comparisons between non-split and DE-split results were not tested within the ANOVA.

Results within the Matrigel-coated TC plastic samples showed that 32.10% ± 6.12% of cells expressed CPM^hi^ fluorescence within the non-split condition, compared to 29.74% ± 6.32% in the DE-split condition (Fig. 3B). For the stiff hydrogel samples, 47.94% ± 8.25% of cells expressed CPM^hi^ fluorescence within the non-split condition versus 33.34% ± 8.23% in the DE- split condition (Fig. 3B). Within soft hydrogel samples, 53.93% ± 7.34% of cells expressed CPM^hi^ fluorescence in the non-split condition, while 31.54% ± 6.72% CPM^hi^ expression was measured in the DE-split condition (Fig. 3B). Comparing the percentage of CPM^hi^ expressing cells, there was a 1.68-fold increase in the soft non-split samples, compared to the Matrigel-coated TC plastic non-split samples (p = 0.0938) (Fig. 3B).

Furthermore, there was a trend of increased CPM^hi^ cells as substrate stiffness decreased among the non-split conditions (Fig. 3B). This same trend did not appear in the DE-split conditions. There were also negligible changes in the percentage of CPM^hi^ expressing cells across all substrate types in the DE-split conditions. These results demonstrate that differentiating cells on soft hydrogels without splitting during the protocol outperformed the outcomes observed with the traditional non-split Matrigel-coated TC plastic approach.

### Elastic Modulus, Cell Line, and Splitting Influence iPSC Differentiation into LP Cells

A DOE statistical approach further analyzed how the input variables of microenvironmental stiffness (elastic modulus), cell line (iPSC Line), and splitting during the differentiation protocol influenced the conversion of iPSCs into LP cells. Combinatorically, these factors have not been systematically studied before, but individually are known to strongly impact lung cell differentiation. The values of 3 x 10^6^ kPa (3 GPa), 18.8 kPa, and 4 kPa corresponded to the stiffness of the Matrigel-coated TC plastic, stiff hydrogels, and soft hydrogels, respectively (Fig. 1B). The three iPSC lines were denoted as lines 1-3 in the DOE. Finally for the splitting variable, the categorical options of non-split or DE-split were included in the DOE numerically as 0 and 1, respectively. All input values were treated as nominal numerical values within the DOE software. Results from IF staining quantification, NKX2.1 gene expression, and CPM^hi^ flow cytometry were added to the DOE model for analysis.

Trend lines for each input variable (Fig. 4A) visually illustrated how response values changed with variations of the input factors. Results revealed that relative NKX2.1 gene expression and the percentage of CPM^hi^ cells were lowest under the 3 GPa microenvironmental stiffness condition, which corresponds to Matrigel on TC plastic (Fig. 4A). Additionally, the DOE analysis indicated that iPSC line 2 cells achieved the highest NKX2.1 relative gene expression and percentage of CPM^hi^ cells (Fig. 4A). Furthermore, splitting at the DE stage was negatively associated with the proportion of cells expressing NKX2.1 protein and CPM^hi^ expression (Fig. 4A). These findings provide invaluable insight for guiding iterative refinements to established differentiation protocols, aiming to improve differentiation efficiencies.

**Figure 4.**
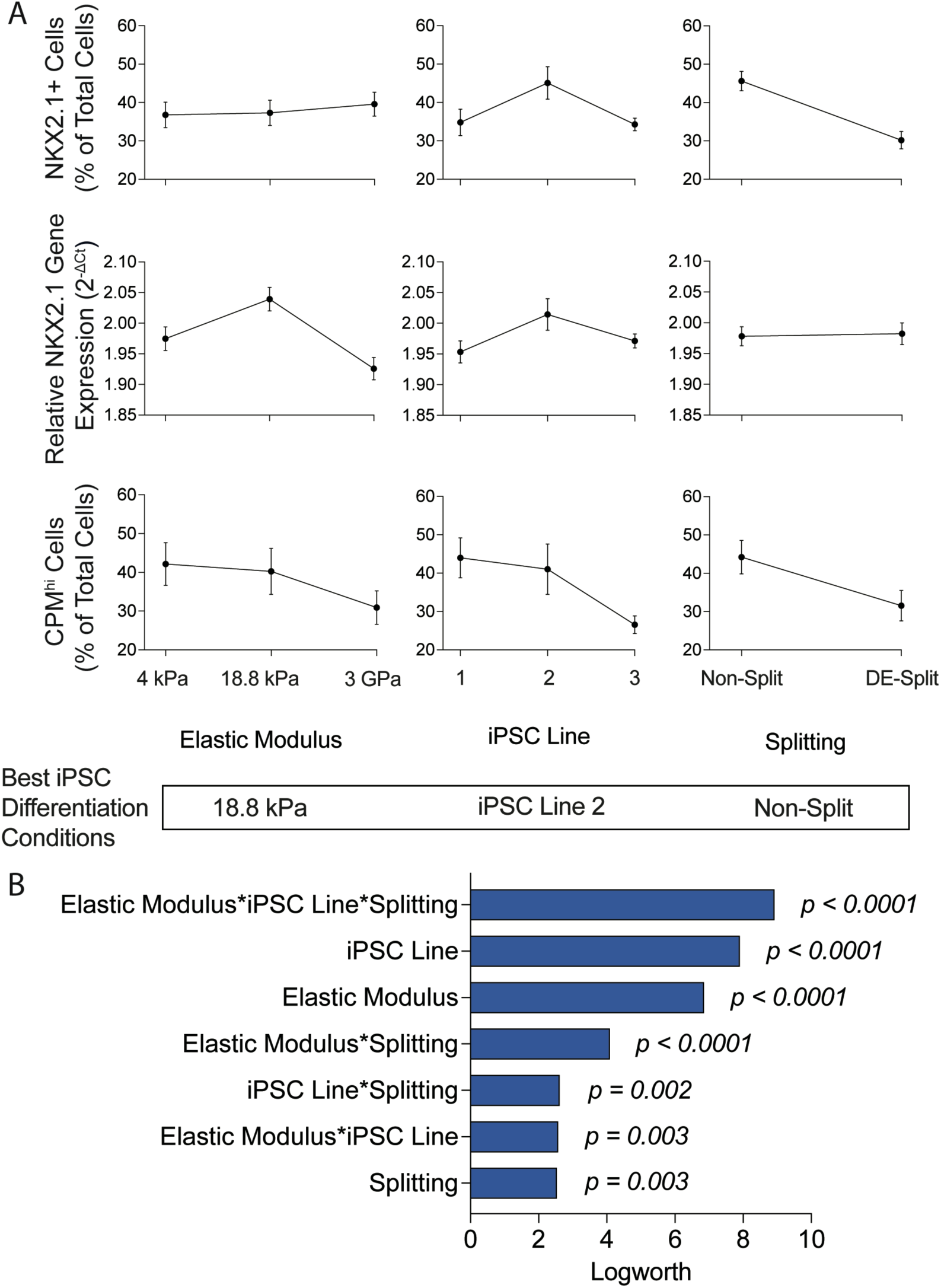
DOE results showed that the elastic modulus, iPSC line, and splitting all significantly influenced LP differentiation outcomes. (A) Trend lines from experimental data showed how the percentage of NKX2.1 cells, NKX2.1 gene expression, and the percentage of CPM^hi^ cells changed in relation to the elastic modulus, iPSC line, and splitting. Data are presented as mean ± SEM. B) The effect magnitude analysis for all factors and interactions showed that the combination of the elastic modulus, iPSC line, and splitting was most influential, and all factors and/or interactions were statistically significant contributors to increasing LP marker expression by the end of the protocol. These results also predicted that the best differentiation conditions would be achieved by using non-split iPSC line 2 cells on stiff hydrogels.

Additionally, the DOE statistically fit model predicted that the highest number of LP cells after 18 days of differentiation would be achieved by using iPSC line 2 on stiff hydrogels that were non-split using the protocol presented here (Fig. 4A). These results also showed that all input variables and interactions among these variables were statistically significant and impacted iPSC differentiation into LP cells. For instance, the interaction effect including elastic modulus, cell line, and splitting had the greatest influence on LP differentiation (p < 0.0001) (Fig. 4B).

Similarly, iPSC line, elastic modulus, and the interaction between elastic modulus and splitting were also significant with p < 0.0001 (Fig. 4B). These results reveal that high levels of variability remain between iPSC cell lines and that substrate elastic modulus is an important consideration when selecting cell culture conditions.

## DISCUSSION

Chronic lung diseases remain the fourth leading cause of death worldwide, with limited treatment options (43). Disease progression leads to impaired stem cell function, tissue damage, and cell death which severely and negatively impact lung health (44, 45). Pulmonary fibrosis is a prominent example, where widespread loss of alveolar epithelial cells significantly diminishes the lung’s capacity for tissue repair and regeneration (19, 46, 47). Similarly, basal stem cell attrition is a key factor underlying several chronic lung diseases (48–51) and is a common consequence of lung transplantation (52–54), contributing to impaired tissue repair and regeneration. To address these challenges, researchers are exploring the therapeutic potential of autologous iPSC-derived progenitor cells, which offer a personalized approach that minimizes immune rejection risk and theoretically supports functional epithelial regeneration. Additionally, an *ex vivo* supply of progenitor cells is crucial for high-throughput studies, such as those focused on optimizing differentiation protocols, testing therapeutic interventions, or *in vitro* disease modeling. Notably, successful transplantation of NKX2.1+ LP-derived basal or alveolar stem cells populations into the airway epithelium have been demonstrated (31, 55–57), underscoring the need for reproducible LP cell production and more efficient, cost-effective differentiation protocols.

This study aimed to minimize reliance on animal-derived products such as Matrigel in LP differentiation protocols by investigating engineered hydrogels as differentiation substrates.

Adopting this engineering approach revealed several factors known to strongly influence cell differentiation, including substrate stiffness (19–21, 58–60), cell line variability (10, 13, 14, 61), and intermediary cell splits (13, 14, 17) , all of which were identified as statistically significant drivers of LP differentiation efficiency. The results demonstrated that tunable stiffness hydrogels were a viable alternative to Matrigel-coated TC-plastic for differentiating iPSCs. Successful LP differentiation on Matrigel-coated TC plastic ranges in efficiency from 5% to 90% NKX2.1- positive or CPM^hi^ cells (62). There were no significant differences in the number of cells expressing NKX2.1 protein between different substrates in the experiments presented here, as measured by IF. However, overall hydrogels outperformed Matrigel-coated TC plastic when gene expression and cell surface markers were evaluated. The highest levels of NKX2.1 gene expression were measured on stiff hydrogels using a non-split protocol and the highest percentage of CPM^hi^ cells was observed in the non-split soft hydrogel condition. Within the non-split samples, the trend of increasing CPM^hi^ cells as substrate stiffness decreased was observed. Based on these results, implementation of a non-split differentiation protocol may be advantageous for certain iPSC lines, consistent with findings from other groups (10, 61).

To minimize the potential confounding variable of sex, three male iPSC lines were used throughout these experiments. These lines were derived from one adult and two neonatal patients. Since iPSC line significantly influenced differentiation results, the pulmonary field will likely benefit from controlling and standardizing iPSC development to reproducibly produce LPs. Future work testing the impact of patient sex and age would also help make these results more representative of all iPSC lines. Although highly controlled, the engineered hydrogels tested in this study differ in composition from the Matrigel-coated TC-plastic samples. This limitation highlights a challenge with Matrigel: its variable composition is difficult to replicate, and it does not allow for precise control over mechanical properties. These differences in composition were not assessed here, representing a potential limitation of this work. Future studies could systematically modify engineered hydrogel composition to better understand how biochemical cues influence iPSC-to-LP differentiation.

Overall, this study pioneers the use of well-defined, tunable-stiffness hydrogels to achieve robust NKX2.1 and CPM^hi^ expression, which successfully marks iPSC differentiation into LP cells. Notably, the elimination of Matrigel represents a key step toward bridging the gap between benchtop research and clinical translation. Unlike existing protocols, this approach offers distinct advantages, including enhanced compliance with regulatory requirements and reduced immunogenic risks, making it more suitable for eventual patient transplantation. The careful design of engineered hydrogel substrates provided physiologically relevant microenvironmental cues, enabling successful differentiation across three independent iPSC lines. This differentiation protocol holds significant promise for broader applications across various cell lines and therapeutic contexts.

## DATA AVAILABILITY

The data that support the findings of this study are openly available in Mendeley Data at doi: 10.17632/cxwk4s76wd.1.

## IRB STATEMENT

All experiments described here used iPSC lines that were derived prior to these studies (Supplemental Table S1) and are classified as human specimens for secondary research. These studies were therefore exempt from Institutional Review Board (IRB) approval.

## DISCLOSURES

C.M.M. is a member of the board of directors for the Colorado BioScience Institute. No conflicts of interest, financial or otherwise, are declared by the other authors.

## AUTHOR CONTRIBUTIONS

Conceptualization, A.E.T., G.B., and C.M.M.; data curation, A.E.T., R.B., and C.M.M; formal analysis, A.E.T., A.L.R., R.B., and C.M.M; project administration, A.E.T. and C.M.M.; supervision, A.L.R. and C.M.M.; writing – original draft, A.E.T. and C.M.M.; writing – review & editing, all authors.

## ACKNOWLEDGEMENTS

The authors would like to acknowledge that Biorender.com was used to create multiple figures for this manuscript. Additionally, the authors would like to thank previous Magin laboratory members Predrag Serbedzija, Donald Campbell, and Jacqueline Miller for helping with initial experiments that motivated this manuscript. We also appreciate Anton Kary (CU Anschutz, Magin laboratory) and Mikala M. Mueller (CU Anschutz, Magin laboratory) for maintaining iPSCs in culture and changing medium that contributed to this work. Shennea McGarvey and the Stem Cell Biobank and Disease Modeling Core (CU Anschutz) also provided significant technical support that helped the Magin laboratory learn iPSC culture techniques in the beginning phases of experiments. Most of the flow cytometric analysis was done in the Barbara Davis Center Bioresource/Molecular Core at the University of Colorado Diabetes Research Center under the supervision and guidance of Scott Beard (CU Anschutz). This core is supported by NIDDK grant # P30-DK116073. FlowJo software was made available for analyzing flow files at the University of Colorado Cancer Center Flow Cytometry Shared Resource Core (CU Anschutz) that is supported by grant P30CA046934. We also would like to acknowledge Dr. Richard Benninger (CU Anschutz, Benninger Imaging and Biophysics Research Lab) and his lab members for offering up their lab space to perform parts of these studies. Dr. Darrell Kotton (Boston University) graciously provided iPSC Line 1 that were used throughout these studies, and we appreciate all the technical support that his laboratory members provided. Furthermore, Dr. Bradford Smith (CU Anschutz, Smith Pulmonary Biomechanics Laboratory) and Anton Kary graciously provided help with statistical data analysis. Lastly, this work was supported by funding from the National Heart, Lung, and Blood Institute of the National Institutes of Health (NIH) under awards R01 HL153096 (CMM, RB, AT) and T32 HL072738 (AT), the National Science Foundation under award 2225554 (CMM, ALR), and the Department of the Army under award W81XWH-20-1-0037 (CMM, AT).

## Notes

https://doi.org/10.17632/cxwk4s76wd.1

