## Supplemental Document for "Engineered hydrogel biomaterials facilitate lung progenitor cell differentiation from induced pluripotent stem cells"

### Materials and Methods

*Maintenance of human induced pluripotent stem cell lines.*

Three human induced pluripotent stem cell (iPSC) (BU3 NGST, FIPS 3c15, and EC11c13, Supplemental Table S1) lines were expanded and maintained on hESC-qualified Matrigel (Corning, cat. # 354277) coated tissue culture (TC) plates for at least two passages before differentiation. More comprehensive characterizations for these iPSC lines can be found in previously published papers (1-6). In brief, to generate the BU3 NGST iPSCs, reprogramming of dermal fibroblasts was performed with the hSTEMCCA-loxP lentiviral vector with four-factors as previously described (7). Similarly, to generate the FIPS 3c15 and EC11c13 iPSCs, reprogramming of human-skin fibroblasts was performed with a polycistronic six-factor lentivirus (1, 5). The iPSCs were maintained in mTeSR Plus (STEMCELL Technologies, cat. # 100-0276) medium with daily medium changes. Confluency was maintained at less than 70-80% during expansion. Passaging was performed as needed using 1X ReLeSR (STEMCELL Technologies, cat. # 05872) (2, 3).

*2D hydrogel synthesis and characterization.*

Circular glass coverslips (18-mm diameter; Electron Microscopy Sciences, cat. #72229-01) were silanized and stored at -20°C for up to 4 weeks before use. In brief, a silanization solution of 90 mL 95% ethanol, 3-4 drops of glacial acetic acid, and 500 µL of (3-Mercaptopropyl)trimethoxysilane (ThermoFisher, cat. #174652500) was prepared in a plastic tricorner beaker and allowed to react while stirring for at least 5 min at room temperature (RT). The coverslips were then passed briefly through a Bunsen burner flame, placed in a Wash-N-Dry coverslip rack (Sigma, cat. #Z743684) and submerged in the silanization solution for 3 min, removed from the silanization solution, rinsed with 95% ethanol, dried at 80°C for 15 min, and finally stored at -20°C.

Glass slides (Corning, cat. #2947-75X50) were treated with Sigmacote (Sigma-Aldrich, cat. #SL2-100ML) just prior to use. 750 mL of Sigmacote was evenly pipetted onto the glass surfaces and left to dry in a chemical fume hood for 20 min at RT. The slides were rinsed with distilled water and dried at 80°C for at least 20 min. Prior to use, the glass slides and coverslips were sprayed with 70% ethanol, transferred into the biosafety cabinet, and left to air-dry.

A poly(ethylene glycol)-norbornene (PEGNB; 8-arm, 10 kg/mol) macromer was synthesized and characterized with nuclear magnetic resonance spectroscopy (1H NMR) using a Bruker DPX-400 FT NMR spectrometer, 300 MHz as previously described (8-10). PEGNB with functionalization of at least 90% based on the comparison of the alkene protons from norbornene to the theoretically expected number of alkene protons, using the ethylene glycol protons as a reference was used in hydrogel formation throughout these studies. Hydrogel formulations contained PEGNB (8-arm, 10 kg/mol), an MMP9-degradable crosslinker (Ac-GCRD-VPLSLYSG-DRCG-NH2, GL Biochem), a laminin peptide mimic (CGYIGSR; 2mM, GL Biochem), a fibronectin peptide mimic (CGRGDS; 2 mM, GL Biochem), laminin/entactin protein (Corning, cat. #354259; 2 mg/mL), and lithium pheyl-2,4,6-trimethylbenzoylphosphinate (LAP; 2.2 mM) photoinitiator. The PEGNB weight percent (wt%) was adjusted to achieve either a soft (5.3 wt%) or stiff (7.75 wt%) final hydrogel. The MMP9-degradable crosslinker was added at a ratio of 0.7 thiol reactive groups to norbornene reactive groups. Stock solutions of the PEGNB, MMP9-degradable crosslinker, RGD, and YIGSR components were individually dissolved in HyClone Dulbecco's Modified Eagle Medium (DMEM)/F12 (Cytiva, cat.

#SH30023.FS). The LAP was protected from light and dissolved in pH 7.2-7.5 (4-(2-hydroxyethyl)-1-piperazineethanesulfonic acid (HEPES, ThermoFisher). To achieve the final concentrations listed above, the respective volumes of each component were combined in a microcentrifuge tube and vortexed.

To create 2D hydrogels, 90 µL of hydrogel precursor solution was pipetted onto Sigmacote-treated glass slides. Then, a functionalized glass coverslip was inverted on top of the hydrogel droplet. Polymerization was initiated by exposure to 365 nm light at 10 $mW/cm^2$ for 5 min. The coverslips with the 0.5 mm thick and covalently bonded hydrogels were transferred into 12 well TC plastic plates (hydrogel side up), submerged in mTeSR Plus medium, and swollen for at least 6 h (37°C, 5% $CO_2$) before cell seeding.

To create hydrogels for rheological characterization, 40 µL of hydrogel precursor solution was pipetted between two Parafilm-covered glass slides that were separated by a 1 mm gasket. This volume was selected to create circular samples of approximately 8-mm diameter. Samples were polymerized by exposure to 365 nm light at 10 $mW/cm^2$ for 5 min. Using a Discovery HR2 rheometer (TA Instruments) and an 8-mm parallel plate geometry, hydrogel mechanics were evaluated. The hydrogels were individually transferred onto a Peltier plate set to 37°C, next the geometry was lowered until the hydrogel was in contact, and a 0.03 N axial force registered on the rheometer. The storage modulus (G') plateau was measured by compressing the hydrogel in increments of 5% until the storage modulus remained constant (11). The plateau for soft hydrogels was 25% compression, and for stiff hydrogels, the plateau was at 30% compression. Hydrogels were then exposed to frequency oscillations with logarithmic sweep with frequencies between 1 and 100 rad $s^{-1}$ at 1% strain. A Poisson's ratio of 0.5 was assumed, and the elastic modulus (E) was calculated assuming the hydrogels exhibited incompressible bulk-elastic characteristics (10, 12-14).

*iPSC to LP cell differentiation.*

Once the undifferentiated iPSCs reached 80% confluency (Day 0), the iPSCs were dissociated for 10 min with 1X Gentle Cell Dissociation Reagent (GCDR, STEMCELL technologies, cat. # 100-0485) and seeded at a density of 200,000 cells/$cm^2$ onto either hESC-qualified Matrigel coated TC-plastic plates, soft hydrogels, or stiff hydrogels. On day 0, the cells were maintained in mTeSR Plus medium supplemented with 10 μM Y-27632 dihydrochloride (Tocris, cat. #1254), a ROCK inhibitor, for 24 h. On day 1, the mTeSR Plus medium was removed manually and replaced with MR+CJ StemDiff definitive endoderm (DE) medium (STEMCELL technologies, cat. #05110) according to the manufacturer's instructions.

On day 2, the medium in each well was manually aspirated and replaced with CJ-only StemDiff DE medium (STEMCELL technologies, cat. #05110) according to the manufacturer's instructions. This process was repeated on days 3 and 4 with CJ-only StemDiff DE medium. On day 5, a subset of samples was split at a 1:3 ratio. For the split samples, the medium was manually removed, and 1 mL of 1X GCDR was added for 10-15 min. The GCDR was then aspirated, and DS/SB medium supplemented with 10 μM Y-27632 dihydrochloride (2, 3) was used to lift the cells off the surfaces. These DE-split cells were then plated back onto the same substrate surface (either freshly prepared Matrigel coated TC-plastic plates or new 2D hydrogels pre-swollen for at least 6 h at 37°C, 5% $CO_2$). Meanwhile, the medium from the wells that contained the non-split samples was removed and replaced with DS/SB medium supplemented with 10 μM Y-27632 (2, 3). On day 6, all samples had medium manually aspirated and DS/SB medium (2, 3) was added. On day 8, the medium was replaced with CBRa medium (2, 3) in each well. The CBRa medium was replenished every 48 h (2, 3) until day 18.

*Immunostaining and image analysis.*

On day 18, cells were dissociated off the cells' respective substrates as described above and cytospun using single cytofunnels (Fisher Scientific, cat. # 59-910-40) to achieve approximately 100,000 cells per slide. Samples were kept hydrated with PBS and then outlined with a hydrophobic pen (Vector Laboratories). Next, the samples were fixed with 4% paraformaldehyde (PFA, Electron Microscopy Sciences) in PBS for 15 min, quenched with 100 mM glycine (Sigma) in PBS for 20 min, and rinsed with PBS for 5 min at RT. Samples were then permeabilized with 0.2% Triton X-100 (Fisher Bioreagents) for 18 min at RT, blocked with 5% bovine serum albumin (BSA, Fisher Scientific) in PBS for 1 h at RT, and incubated overnight at 4 °C with the rabbit anti-human NK2 homeobox 1 (NKX2.1) antibody (R&D Systems, cat. #MAB94581) at a final concentration of 20 µg/mL in immunofluorescence (IF) solution that contained 3% BSA and 0.1% Tween 20 (Sigma) in PBS. After overnight primary antibody incubation, samples were rinsed three times in IF solution and incubated with the secondary donkey anti-rabbit Alexa Fluor 488 antibody (Jackson ImmunoResearch, cat. #711-545-152) in IF solution for 45 min at RT using a 1:300 dilution and while protected from light. Samples were washed three times in IF solution and then incubated with 4',6-diamidino-2-phenylindole (DAPI) (Sigma-Aldrich, cat. #D9542) diluted 1:1000 in PBS for 15 min at RT, protected from light. Finally, samples were rinsed with deionized water three times, mounted with Prolong Gold Antifade Reagent (Fisher Scientific, cat. #P36930), cover-slipped, and stored protected from light until imaging. Negative controls were prepared identically, except the primary antibody was omitted.

Cytospun samples that were IF-stained were imaged on an Olympus BX63 upright microscope. Six random areas per slide were imaged with DAPI and FITC filters under a 4x objective. Images were analyzed with ImageJ Software (NIH). Background was subtracted from images across all three cell lines based on representative negative control images. The ImageJ plugin StarDist quantified the number of NKX2.1-positive cells and the total number of cells present. The versatile (fluorescent nuclei) model was used with the percentile range from 1 to 100, both the probability/score threshold and overlap threshold values were set to 0.4, and all other parameters remained as the default values. To determine the final percentage of cells positive for NKX2.1 staining, individual NKX2.1-positive cell number readings were normalized against respective total number of cells present.

*RNA Isolation and cDNA Synthesis, and gene expression analysis.*

On day 18, after cells were lifted off the respective substrates, cells were centrifuged at 300 x g for 5 min at 4 °C. Next, the supernatant medium was aspirated, and the cell pellet was resuspended in 700 µL of TRIzol Reagent (Fisher Scientific, cat. #15-596-026). These samples were pipetted and briefly vortexed, left for at least 5 min at RT and then stored at -80 °C for up to 4 weeks. Following, samples were thawed and 100 µL of 1-bromo-3-chloropropane (BCP; Fisher Scientific) was added to each sample. The samples were vortexed, incubated at RT for 5 min, and then incubated on ice for an additional 5 min. The samples were then centrifuged at 12,000 x g for 15 min so the clear layer of the sample could be transferred into a separate RNAse-free 1.5 mL microcentrifuge tube. An equal amount of 100% ethanol (EtOH) was added to the clear RNA layer volume and the two were briefly vortexed. Up to 700 µL of total volume was transferred into RNeasy Mini Kit columns (Qiagen, cat. #74104). The tubes were centrifuged at 8,000 x g for 30 s. The flow through was discarded and then 700 µL of RW1 buffer (Qiagen) provided by the kit was added to the column. The samples were then centrifuged at 8,000 x g for 30 s and the flow through was discarded again. Next, 500 μL of RPE buffer (Qiagen) was added to the column, centrifuged at 8,000 x g for 30 s, and the flow through was discarded. Again, 500 μL of RPE buffer (Qiagen) was added to the column, centrifuged at 8,000 x g for 30 s, and the flow through was discarded. To dry the samples completely, the columns were transferred into new 1.5 mL provided tubes and spun at 17,000 x g for 2 min. Finally, 30 μL of RNAse free water (Qiagen) was added directly on top of the membrane and one last centrifuge step was completed at 17,000 x g for 1 min. RNA quantity and purity, as assessed by the ratio of absorbance readings taken at 260 nm and 280 nm ($A_{260}/A_{280}$), were measured using a BioTek plate reader and a Take3 Micro-Volume Plate. The isolated RNA was then converted into cDNA using a high-capacity cDNA Reverse Transcription Kit (Applied Biosystems, cat. #4368814) according to the manufacturer's protocol.

*Assessment of Gene Expression.*

Reverse transcription-quantitative polymerase chain reaction (RT-qPCR) assessed gene expression of NKX2.1 in all sample conditions and was normalized to housekeeping gene ribosomal protein S18 (RPS18). iTaq Universal SYBR Green Supermix (Bio-Rad, cat. #1725121) and a CFX Opus 96 (Bio-Rad) were used for all experiments. All gene expression data underwent a natural log transformation to normalize the data before relative gene expression was calculated using a $2^{-\Delta Ct}$ approach. After statistical analyses were performed, values were untransformed (by taking the exponent of the natural logged value) and then presented in figures.

*Flow cytometry.*

Cells collected at the end of lung progenitor induction (day 18) were dissociated from samples using 1X GCDR. GCDR was left on the cells for either 10 min (hESC-qualified

Matrigel coated TC-plastic samples) or 15 min (soft and stiff hydrogel samples). Manually the GCDR was aspirated, and using a pipette tip, cross-hatching (scratching lines across the material surface) was performed to help facilitate dissociation. All cells were collected and DMEM/F12 was used to dilute the GCDR. The cells were gently pipetted to ensure a single cell suspension was achieved, then passed through a 70 μm strainer (CELLTREAT, cat. #229483) and resuspended in flow cytometry (FC) buffer containing 2% fetal bovine serum (FBS, Cytiva, cat. #SH30071.03) in phosphate buffered saline (PBS, Cytiva, cat. #SH30256.FS). After counting, a proportion of cells were kept in FC buffer at 4 °C for 15 min, and used as unstained and secondary-only controls (~1 x $10^5$ cells in each). The remaining cells were stained for 15 min at 4 °C with a mouse anti-human carboxypeptidase M (CPM, Fujifilm Wako Chemicals, cat. #014-27501) at a concentration of 0.5 μL CPM antibody for each 100,000 cells/100 μL FC buffer. After incubation, 1 mL of PBS was added to all test tubes to help wash the cells. The tubes were then centrifuged at 300 x g for 5 min at 4 °C to pellet the cells.

Next, the secondary-only control and cells from each respective sample condition were stained for 30 min at 4 °C with a rat anti-mouse PE-Cyanine7 antibody (PE-Cy7, Thermo Scientific, cat. # 25-4015-82) at a concentration of 2.5 μL PE-Cy7 antibody for each 1 × $10^6$ cells/100 μL FC buffer. Simultaneously, the unstained controls underwent the same treatment but excluded the secondary antibody. Starting with the secondary antibody staining step, samples were protected from light. After incubation, 1 mL of PBS was added to all the test tubes to help wash the cells and dilute out any antibody present. The test tubes were then centrifuged at 300 x g for 5 min at 4 °C. To stain for viability, Calcein Violet-AM (Fisher Scientific, cat. #501145046) was added to FC buffer so the final concentration was 0.01 μM. For samples requiring Calcein Violet-AM staining, the cells were resuspended in this staining solution and incubated for at least 15 min at 4 °C. For all other samples, the cells were resuspended and maintained in FC buffer.

To prepare the unstained beads and beads + PE-Cy7 controls, Ultracomp eBeads (Thermo Scientific, cat. #01-2222-42) were used. For each control, 3 drops of eBeads were added to a flow test tube. For the beads +PE-Cy7 control, 2 μL of PE-Cy7 antibody was also added. Finally, following the last incubation, flow cytometric analysis was completed using all prepared controls and samples. While compensating on the machine, eFluor 450 served as a sufficient substitute for Calcein Violet-AM as eFluor 450 was the available fluorophore option on the spectral cytometer. eFluor 450 has a maximum excitation/emission spectra of 405/450 nm, while Calcein Violet AM has a maximum excitation/emission spectra of 400/452 nm.

In total, the following samples were prepared to provide sufficient controls and assist in selecting gates for flow cytometric analysis: 1) unstained Ultracomp eBeads 2) Ultracomp eBeads + PE-Cy7, 3) unstained and undifferentiated (day 0) iPSCs, 4) undifferentiated (day 0) iPSCs with Calcein Violet-AM, 5) unstained day 18 cells, 6) day 18 cells stained with only PE-Cy7, 7) day 18 non-split cells from Matrigel coated TC-plastic stained with CPM + PE-Cy7+ Calcein Violet-AM, 8) day 18 DE-split cells from Matrigel coated TC-plastic stained with CPM + PE-Cy7+ Calcein Violet-AM, 9) day 18 non-split cells from soft hydrogels stained with CPM + PE-Cy7+ Calcein Violet-AM, 10) day 18 DE-split cells from soft hydrogels stained with CPM + PE-Cy7+ Calcein Violet-AM, 11) day 18 non-split cells from stiff hydrogels stained with CPM + PE-Cy7+ Calcein Violet-AM, and 12) day 18 DE-split cells from stiff hydrogels stained with CPM + PE-Cy7+ Calcein Violet-AM. All flow cytometric analysis was performed on a CYTEK Aurora 5 laser spectral cytometer (CU Anschutz Barbara Davis Bioresource/Molecular Core).

*Statistical Analysis.*

For all IF and RT-qPCR quantification, results included three biological replicates with six to eight technical replicates. For flow cytometry quantification, results included three biological replicates with three to four technical replicates. RT-qPCR results were normalized using a natural log transformation. Meanwhile, the IF and flow datasets were assessed for normality using a Shapiro-Wilk test and passed. Two-way ANOVA and a Tukey test for multiple comparisons assessed statistical differences within non-split or DE split conditions for immunofluorescence, gene expression, and flow datasets (GraphPad Prism). To identify the best predicted differentiation conditions, the influence of substrate elastic modulus, iPSC Line, and splitting on the total number of $CPM^{hi}$ cells, NKX2.1 gene expression, and the percentage of NKX2.1+ cells were investigated with a design of experiments (DOE) approach using JMP software (Pro 18 Version, SAS). The measured percentage of $CPM^{hi}$ cells, NKX2.1 gene expression, and percentage of NKX2.1 cells were entered back into the DOE and a standard least squares model was applied. Statistical analysis was performed using a three-way ANOVA with all interactions terms to fit the model and identify the best fit lines. Statistical significance was defined as having a P value <0.05.

**Results**

**Table S1**. iPSC line demographics.

| iPSC Line | Technical Name | Sex | Age (Years) | Race/ Ethnicity | Tissue Origin | Lab Origin |
|---|---|---|---|---|---|---|
| 1 | BU3 NGST | Male | 32 | Caucasian | Healthy/ Normal Skin | Boston University, CReM Institute; Darrell Kotton Lab |
| 2 | FIPS 3c15 | Male | Neonatal | Unknown | Healthy/ Normal Foreskin | University of Iowa; Amy Ryan Lab |
| 3 | EC11c13 | Male | Neonatal | Unknown | Cord Blood | University of Iowa; Amy Ryan Lab |

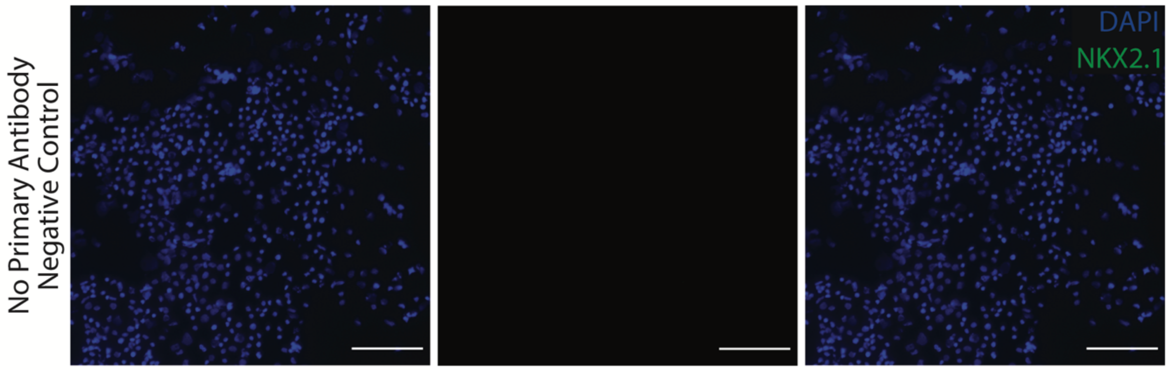


**Fig. S1**. Representative immunofluorescence staining of a negative control (no primary antibody) sample confirmed no unspecific secondary staining occurred. These images were obtained from day 18 cytospun cells (iPSC Line 3) that underwent a DE-split, were stained for NKX2.1 (green), and counterstained with DAPI (blue). Scale bar = 100 $\mu$m.
